# ReviewBench: An Extensible Framework for Benchmarking Human and AI Manuscript Review

**DOI:** 10.64898/2026.04.17.719279

**Authors:** Natalie N Khalil, TJ Reed, Matteo R Ciccozzi

## Abstract

The volume of scientific manuscripts is rising faster than the available pool of expert reviewers, and AI tools are emerging as a possible response, ranging from frontier large language models applied directly to peer review to purpose-built multi-agent systems. Scalable, standardized benchmarks are needed to regularly evaluate how these tools compare to one another and to human reviewers. We present ReviewBench, an open-source, venue-agnostic framework that compares human and AI reviews across structure, alignment with a paper’s major claims, impact, and critique category. We apply ReviewBench to 145,021 review comments from human reviewers, frontier large language models (GPT-5.2, and Gemini 3 Pro), and Reviewer3.com (R3), a multi-agent peer review system. The dataset spans papers in computer science (ICLR 2025, *n* = 1,000), social science (Nature Human Behaviour, *n* = 142), and life science (eLife, *n* = 1,000). Across disciplines, AI reviews are more structured and engage more directly with a paper’s major claims, with R3 more often surfacing consequential comments, defined as comments capable of undermining those claims. When restricting to critical comments, however, human reviewers rank first on consequential rate on more individual papers than any AI source, despite a lower average. We identify a bimodal reviewer distribution with peaks near 0% and 100%, indicating that many reviewers outperform AI on this metric, but a substantial fraction of reviewers near 0% brings the average down. Critique typing demonstrates systematic differences, where humans emphasize contribution and clarity, while AI emphasizes validity, sufficiency, and transparency. Together, these findings argue against framing AI as a replacement for human review and instead support a complementary model in which AI scales technical verification of major claims while human judgment remains essential for evaluating contribution and shaping editorial decisions.

## 1 Introduction

Scientific publications are growing rapidly. The total number of articles indexed in Scopus and Web of Science was roughly 47% higher in 2022 than in 2016, following an exponential trajectory of approximately 5.6% per year [1]. Artificial intelligence (AI) tools accelerate this by lowering barriers to manuscript production, ranging from support for literature review [2, 3, 4] and writing [5] to increasingly autonomous scientific systems [6, 7]. Researchers who incorporate AI methods into their work publish 3.02 times more papers [8].

Peer review has not kept pace. Across multiple disciplines, reviewer acceptance rates are declining [9], the total volume of invitations is increasing [10, 11], and the median time from submission to final peer-reviewed decision exceeds six months [12]. In a 2016 Publons survey, 75% of editors identified “finding reviewers and getting them to accept review invitations” as the most difficult part of their work [10]. The reviewer pool is not growing proportionally with submission volume, creating a fundamental sustainability problem for publishing.

AI tools for manuscript review have also emerged [13, 14, 15, 16, 17], ranging from frontier large language models (LLMs) that have already been applied directly to peer review [18] to tool-augmented multi-agent systems with search capabilities [14, 19], such as Reviewer3.com (R3). As models improve and AI review initiatives multiply, it is unclear how these tools comparatively perform. Automated, scalable benchmarks are needed to evaluate them.

Because reviews are unstructured, comparing them requires a shared reference frame. Recent work converts papers into structured claim representations [20, 21], in some cases grouping atomic claims into results and comparing human and machine evaluations at the result level [21]. We extend this approach by parsing reviews into individual comments, extracting a paper’s major claims, and mapping each human and AI comment to this fixed reference frame. We then assign a stance (critical, supportive, or neutral) for that claim. Because comments sharing the same claim and stance can vary in impact, we add a consequential label that distinguishes challenges capable of undermining the mapped claim from minor concerns. Each comment is further decomposed into its structural elements, including the specific issue identified, its justification, a suggested remedy, and an anchor to a specific location in the paper, and assigned a critique category (validity, sufficiency, contribution, clarity, or transparency). Together, these criteria enable direct comparison of human and AI reviews across structure, claim alignment, and critique category.

We present ReviewBench, an open-source evaluation framework that is extensible across AI review systems and publication venues with openly available human reviews. We apply ReviewBench to 145,021 review comments from humans, R3, and frontier large language models GPT-5.2 and Gemini 3 Pro on computer science (ICLR 2025, *n* = 1,000), social science (Nature Human Behaviour, *n* = 142), and life science (eLife, *n* = 1,000) papers. Across disciplines, AI reviews are more structured than human reviews and engage more directly with a paper’s major claims, with R3 most often surfacing comments capable of undermining those claims. When restricting to critical comments, human reviewers rank first on consequential rate on more papers than any AI source, despite a lower average. We identify a bimodal reviewer distribution with peaks near 0% and 100%, indicating that the most consequential human reviewers outperform AI on this metric, but a substantial fraction of reviewers near 0% brings the average down. Human and AI sources also differ systematically in critique focus, with humans emphasizing contribution and clarity, while AI focuses on validity, sufficiency, and transparency. Together, these findings point to a complementary role for AI in scaling technical verification of major claims while human judgment remains essential for evaluating contribution and shaping editorial decisions.

## 2 Results

### 2.1 Overview

ReviewBench is an extensible benchmark framework consisting of raw data fetchers, a canonicalization layer, AI review generators, and a processor for assessment and scoring (Figure S1). We apply ReviewBench to computer science (ICLR 2025, *n* = 1,000), social science (Nature Human Behaviour, *n* = 142), and life science (eLife, *n* = 1,000) papers (Figure S2A). The ICLR dataset includes 288 accepted and 712 rejected papers (28.8% acceptance rate) (Figure S2B), spanning 21 primary topic areas, with the largest concentrations in ‘foundation or frontier models’ (Figure S2C). The Nature Human Behaviour dataset spans article keywords including ‘science, technology, and society’, ‘economics’, and ‘cognitive neuroscience’ (Figure S2D). eLife data was loaded without topic metadata [22]. No filtering was applied based on topic or acceptance decision and the ICLR sample’s acceptance rate closely approximates the overall ICLR 2025 acceptance rate of 32% [23].

To assess whether evaluator choice would materially affect conclusions, GPT-5.2 and Gemini 3 Pro independently assessed the same comments on 50 randomly sampled papers (Figure S3). The Gemini evaluator, used throughout this paper, shows no self-favoritism on any assessment metric used in this study (Figure S4). Both evaluators rank sources (human, R3, GPT-5.2, and Gemini 3 Pro) similarly, agreeing on the first-place source for all metrics and on exact rank ordering for all but one metric (Supplementary Table S1). Differences therefore reflect general calibration rather than source-specific bias.

### 2.2 AI Reviews Are More Structured

Across three venues, the pipeline assessed 145,021 review comments from human reviewers, R3, and frontier large language models GPT-5.2 and Gemini 3 Pro (Figure 1A). Human comment volume and length vary by venue, with ICLR having the highest comment count per paper but the shortest comments, and Nature Human Behaviour having fewer but longer comments, reflecting differences in venue review format (Figure 1B–C). In contrast, the AI sources’ number of comments and comment lengths are stable across venues. GPT-5.2 produces the most comments of all AI sources across all three venues with large effect sizes (all *p*_adj_ < .001, |*d*| ≥ 1.09), followed by R3, then Gemini 3 Pro.

**Figure 1:**
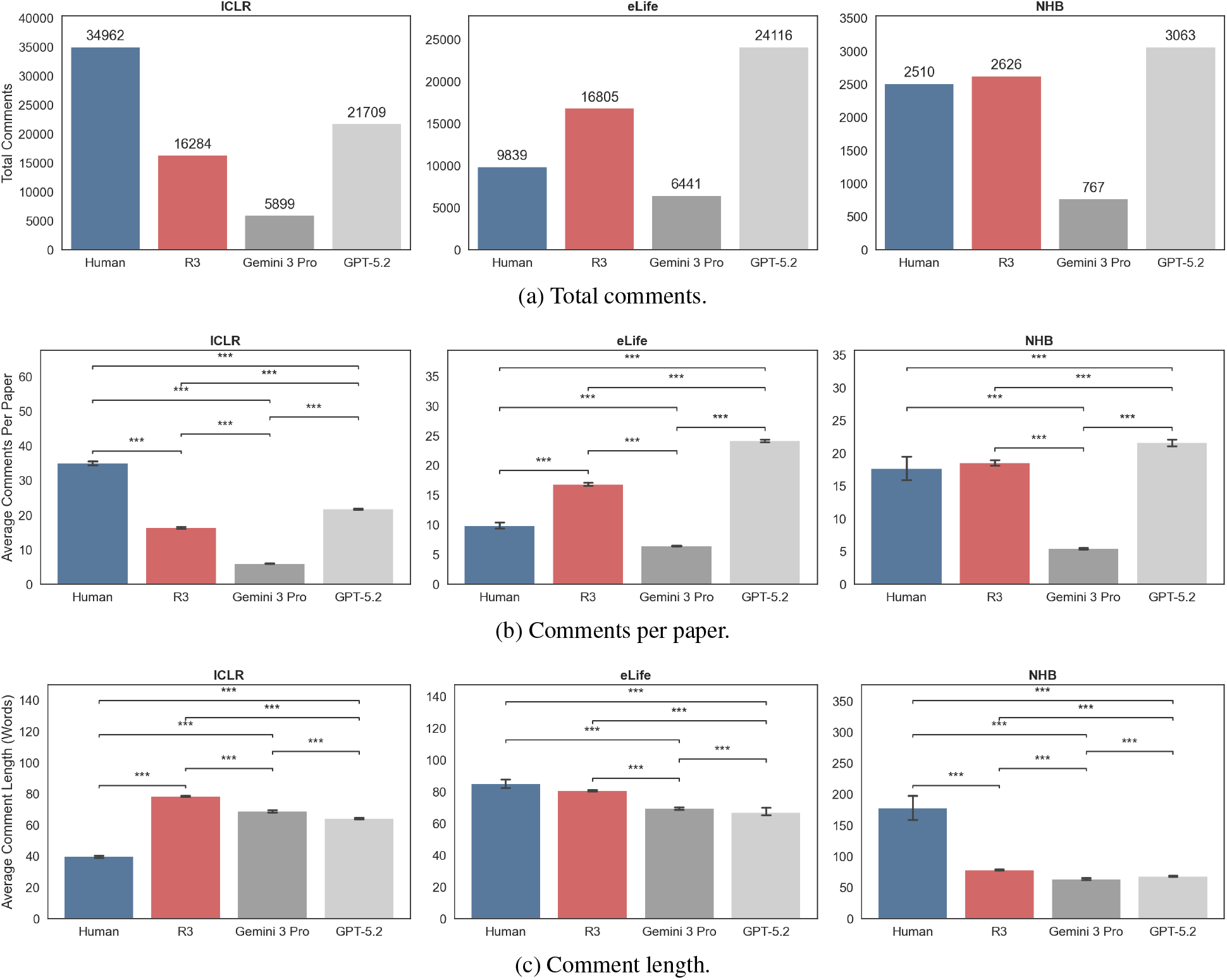
Overview. Total comments, comments per paper, and comment length by source across three venues. NHB = Nature Human Behaviour. Bars show per-paper means, error bars show 95% CIs. Brackets indicate significance (* *p* < .05, * \**p* < .01, * * \**p* < .001, Wilcoxon signed-rank, Holm–Bonferroni corrected). Cohen’s *d* reported per venue (ICLR / eLife / NHB) for pairs: Human–R3, Human–Gemini, Human–GPT, R3–Gemini, R3–GPT, Gemini– GPT. (a) Total comment count by source. Aggregate counts. (b) Mean comments per paper. ICLR *n* = 1,000, eLife *n* = 1,000, NHB *n* = 142. *d*: 2.37, 3.97, 1.74, 3.30, −1.42, −6.52 / −1.15, 0.64, −2.37, 3.35, −1.73, −5.87 / −0.10, 1.57, −0.48, 7.12, −1.09, −7.32. (c) Average comment length (words). ICLR *n* = 1,000, eLife *n* = 999–1,000, NHB *n* = 142. *d*: −4.40, −2.61, −2.59, 0.93, 1.69, 0.42 / 0.14, 0.49, 0.43, 1.08, 0.47, 0.09 / 1.18, 1.35, 1.30, 1.97, 1.54, −0.54.

Peer review comment structure can include the specific issue identified, the justification for why it matters, how to fix it, and where it exists in the text. We extracted these items verbatim from all comments and calculated per-paper specification rate, justification rate, actionability rate, and anchored rate, respectively. AI reviews are more structured than human reviews across all three disciplines, with R3 leading on justification (96.4–97.3%, all *p*_adj_ < .001, |*d*| = 1.74 vs. human) and actionability (99.2–99.9%, all *p*_adj_ < .001, |*d*| = 1.38 vs. human) (Figure 2).

**Figure 2:**
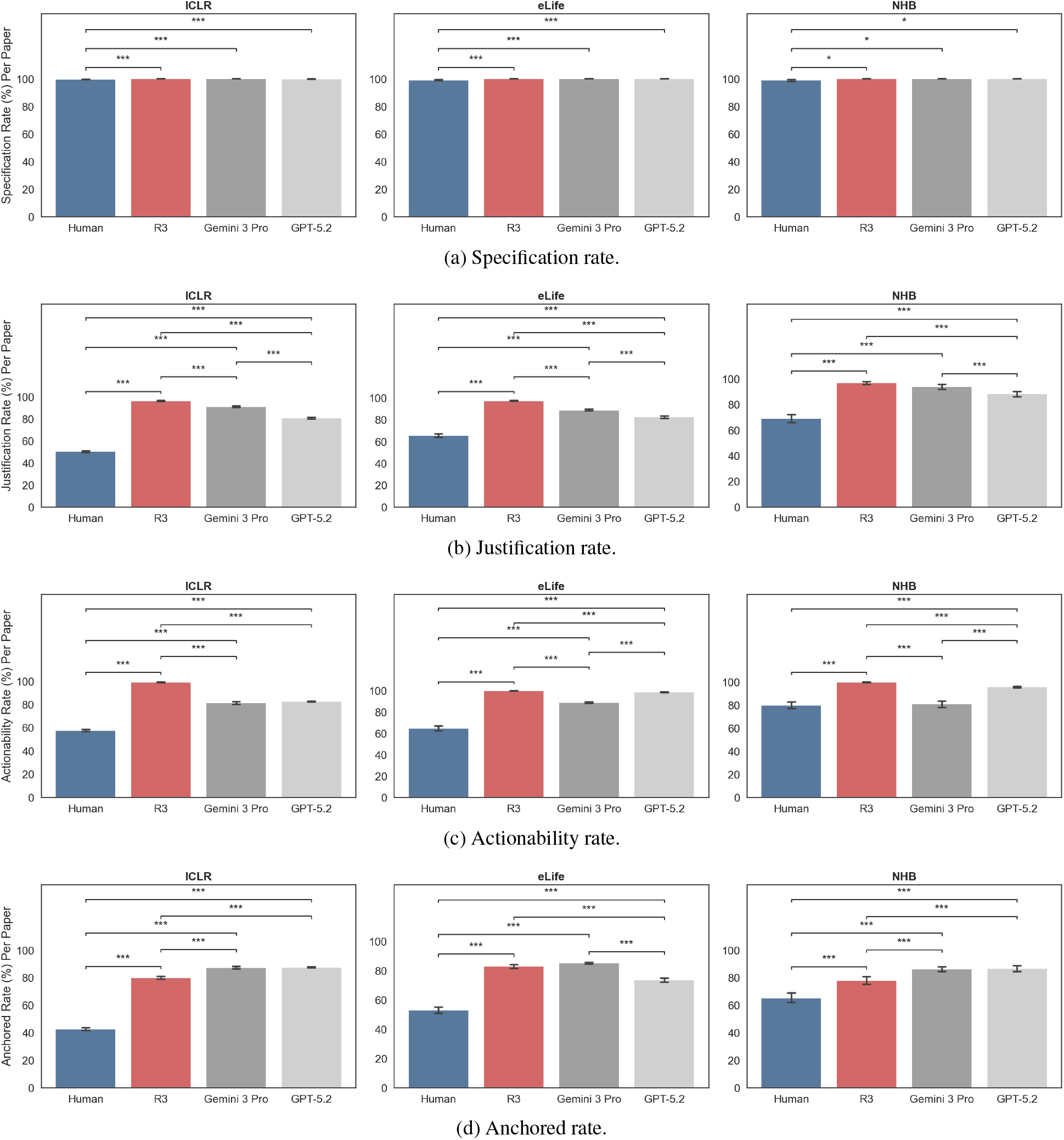
Comment Structure. Specification, justification, actionability, and anchored rates by source across three venues. NHB = Nature Human Behaviour. Bars show per-paper means, error bars show 95% CIs. Brackets indicate significance (\**p* < .05, * * *p* < .01, * * \**p* < .001, Wilcoxon signed-rank, Holm–Bonferroni corrected). Cohen’s *d* reported per venue (ICLR / eLife / NHB) for pairs: Human–R3, Human–Gemini, Human–GPT, R3–Gemini, R3– GPT, Gemini–GPT. ICLR *n* = 1,000, eLife *n* = 999–1,000, NHB *n* = 142. (a) Specification rate: percentage of comments that identify a specific issue. *d*: −0.28, −0.28, −0.21, 0.00, 0.12, 0.12 / −0.21, −0.21, −0.20, 0.00, 0.06, 0.06 / −0.31, −0.31, −0.31, 0.00, 0.00, 0.00. (b) Justification rate: percentage of comments that explain why the issue matters. *d*: −4.06, −3.10, −2.16, 0.52, 1.40, 0.80 / −1.74, −1.17, −0.80, 0.78, 1.19, 0.43 / −1.89, −1.54,−1.19, 0.31, 0.93, 0.50. (c) Actionability rate: percentage of comments that include a suggested remedy. *d*: −3.98, −1.37, −2.27, 1.26, 3.59, −0.08 / −1.38, −0.92, −1.32, 1.70, 0.45, −1.38 / −1.53, −0.06, −1.19, 1.68, 1.46, −1.28. (d)Anchored rate: percentage of comments that reference a specific location in the paper. *d*: −2.29, −2.91, −3.47, −0.49, −0.61, −0.03 / −1.09, −1.29, −0.70, −0.15, 0.43, 0.63 / −0.66, −1.25, −1.24, −0.59, −0.60, −0.04.

### 2.3 AI Sources Engage Directly With Major Claims

For each paper, the pipeline extracted 3–7 major claims (10,388 total) and verbatim source excerpts (Figure S5). The pipeline then maps review comments to the most relevant claim ID, or unmapped if overly broad (spanning multiple claims) or editorial (spanning no claims). R3 (63.5–79.0%) and Gemini 3 Pro (66.3–78.0%) achieve higher mapping rates than all other sources across venues (all *p*_adj_ < .001, |*d*≥ | 0.48) (Figure 3A), suggesting that these sources may engage more directly with major claims.

**Figure 3:**
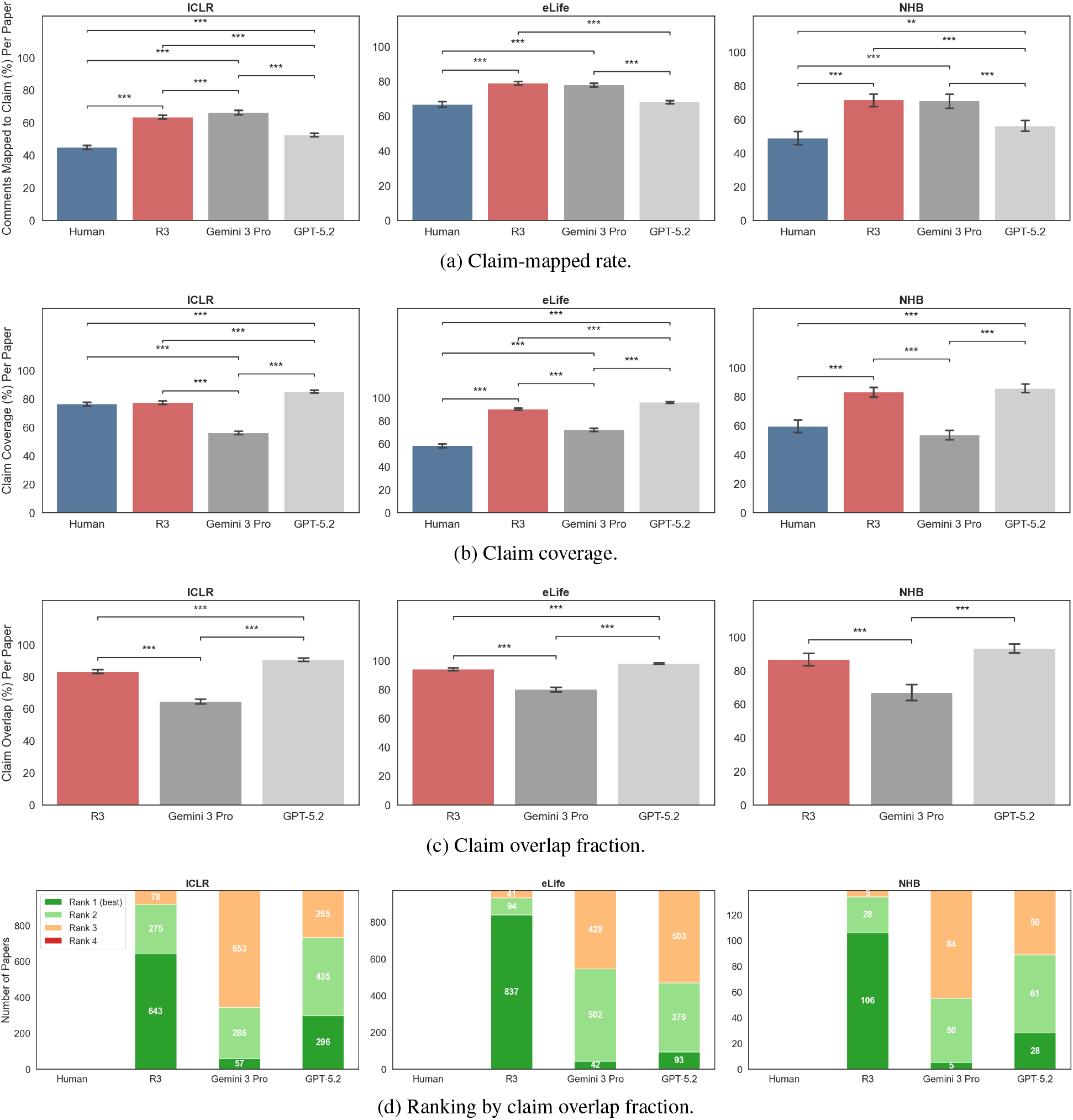
Claim Mapping. Claim-mapped rate, claim coverage, claim overlap fraction, and per-paper ranking by source across three venues. NHB = Nature Human Behaviour. Bars show per-paper means, error bars show 95% CIs. Brackets indicate significance (\**p* < .05, * \**p* < .01, * * \**p* < .001, Wilcoxon signed-rank, Holm–Bonferroni corrected). Cohen’s *d* reported per venue (ICLR / eLife / NHB) for pairs: Human–R3, Human–Gemini, Human–GPT, R3–Gemini, R3–GPT, Gemini–GPT (panels a–b) or R3–Gemini, R3–GPT, Gemini–GPT (panel c). (a) Claim-mapped rate: percentage of comments mapped to a specific paper claim. ICLR *n* = 1,000, eLife *n* = 999–1,000, NHB *n* = 142. *d*: − 0.96, − 1.01, − 0.41, − 0.13, 0.58, 0.66 / − 0.56, − 0.48, − 0.06, 0.06, 0.74, 0.60 / − 0.94, − 0.86, − 0.31, 0.02, 0.70, 0.62. (b) Claim coverage: percentage of each paper’s extracted claims addressed by at least one comment. ICLR *n* = 1,000, eLife *n* = 1,000, NHB *n* = 142. *d*: − 0.05, 0.97, −0.47, 1.08, −0.45, −1.59 / −1.50, −0.61,−1.90, 1.05, −0.47, −1.56 /−1.00, 0.25, −1.14, 1.51, −0.14, −1.71. (c) Claim overlap fraction: percentage of human-addressed claims also addressed by each AI source. ICLR *n* = 996, eLife *n* = 972, NHB *n* = 139. *d*: 0.86, 0.42, 1.29 / 0.68, 0.30, 0.94 / 0.76, 0.34, 1.14. (d) Per-paper ranking by claim overlap fraction. For each paper, AI sources are ranked from highest to lowest claim overlap fraction with human reviewers.

Claim coverage measures the fraction of a paper’s extracted claims addressed by at least one comment (Figure 3B). GPT-5.2 leads on claim coverage across all three venues (85.4–96.2%, all *p*_adj_ < .001, |*d*|≥ 0.45 except GPT-5.2 vs. R3 in Nature Human Behaviour where *p*_adj_ = 1.0), consistent with its larger comment volume. In ICLR, R3 and human reviewers have comparable claim coverage (77.5% vs. 76.5%, *p*_adj_ = 1.0). Gemini 3 Pro has the lowest claim coverage of all AI sources, consistent with its smaller comment volume.

We also measure the claim overlap fraction, defined as the percentage of human-addressed claims also addressed by each AI source (Figure 3C). GPT-5.2 achieves the highest average overlap with human reviewers across all three venues (90.6–98.0%, all *p*_adj_ < .001, |*d*| ≥ 0.30 except GPT-5.2 vs. R3 in Nature Human Behaviour where *p*_adj_ = 0.052), again consistent with its greater comment volume, but per-paper rankings show that R3 ranks first on claim overlap on more individual papers in every venue (Figure 3D).

### 2.4 Human Reviews Have a Balanced Stance Profile

The pipeline assigns a stance (supportive, critical, or neutral) to mapped comments (for examples, see Supplementary Table S2). We calculate stance rates as aggregate proportions of each stance over all comments mapped to a claim (Figure 4A). Across all venues, humans have the most balanced stance profile, with the highest supportive (11.1– 26.8%) and neutral (8.0–8.2%) rates and the lowest critical rates (65.1–80.9%) in most venues. R3 is the most critical of all sources (98.0–98.3%) across all three venues. Gemini 3 Pro and GPT-5.2 are intermediate in all three stance categories.

**Figure 4:**
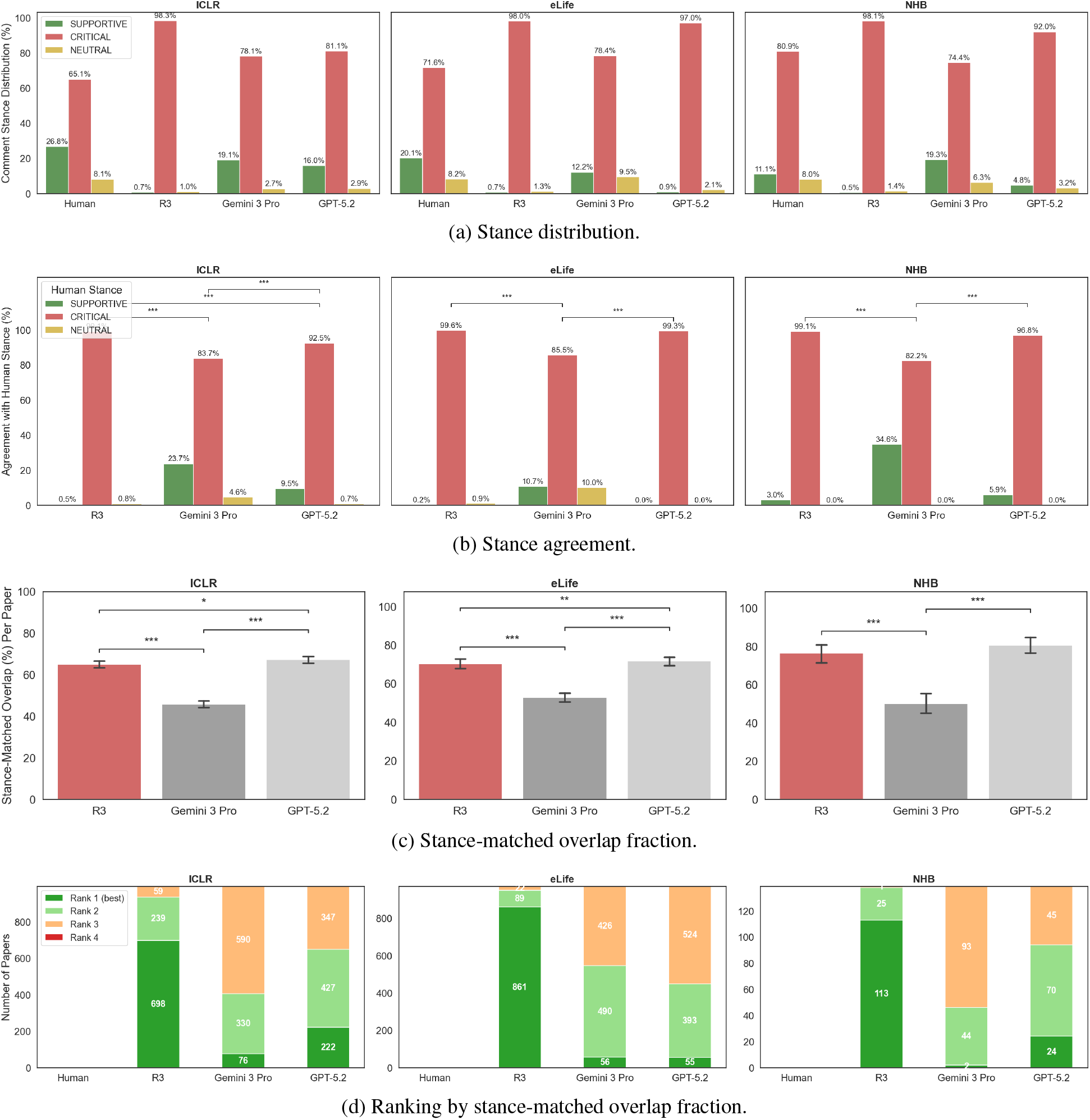
Stance Distribution and Agreement. Stance distribution, stance agreement, stance-matched overlap fraction, and per-paper ranking by source across three venues. NHB = Nature Human Behaviour. Bars show per-paper means, error bars show 95% CIs. Brackets indicate significance (\**p* < .05, * \**p* < .01, * * \**p* < .001, Wilcoxon signed-rank, Holm–Bonferroni corrected). Cohen’s *d* reported per venue (ICLR / eLife / NHB) for pairs: R3–Gemini, R3–GPT, Gemini–GPT (panels b–c). (a) Stance distribution: proportion of claim-mapped comments classified as supportive, critical, or neutral. Aggregate percentages. (b) Stance agreement: for each stance category, the fraction of shared claims where the AI source assigns the same stance as the human. ICLR *n* = 916–948, eLife *n* = 776–808, NHB *n* = 124–131. *d* (critical): 0.67, 0.49, −0.36 / 0.67, 0.02, −0.68 / 0.73, 0.23, −0.58. (c) Stance-matched overlap fraction: percentage of all human-addressed claims that the AI source also addresses with the same dominant stance. ICLR *n* = 996, eLife *n* = 972, NHB *n* = 139. *d*: 0.72, − 0.09, − 0.82 / 0.47, − 0.04, −0.51 / 0.88, −0.16, − 1.08.(d) Per-paper ranking by stance-matched overlap fraction. For each paper, AI sources are ranked from highest to lowest.

For each stance category, we measure stance alignment as the fraction of shared claims where the AI source assigns the same stance as the human (Figure 4B). R3 agrees with human critical judgments most frequently, but Cohen’s *κ* confirms this is at chance level (*κ* = 0.001–0.057) given R3’s near-universal critical stance.

We also compute a stance-matched overlap fraction as the fraction of all human-addressed claims that the AI source also addresses with the same dominant stance (Figure 4C). R3 and GPT-5.2 achieve comparable stance-matched overlap fraction (64.8–76.3% and 67.1–80.7%, respectively) with negligible effect sizes (*d* < 0.2), while Gemini 3 Pro is substantially lower (45.7–52.7%). Per-paper rankings again show R3 ranks first most frequently on stance-matched overlap across all three venues (Figure 4D).

### 2.5 R3 Comments Are More Frequently Consequential to Major Claims

Comments can map to the same claim and stance but vary in impact. To differentiate these, the pipeline assigns each mapped comment a binary consequential label. Consequential comments are defined as comments that could identify an issue that undermines the mapped claim (for examples, see Supplementary Table S3). Consequential rate is the per-paper percentage of comments mapping to a claim ID that also have a consequential label.

R3 achieves the highest consequential rate across all three venues (87.9–93.2%), compared to 65.5–77.5% for Gemini 3 Pro (all *p*_adj_ < .001, |*d*| ≥ 0.79), 73.7–81.4% for GPT-5.2 (all *p*_adj_ < .001, |*d*| ≥ 0.50), and 60.0–67.7% for humans (all *p*_adj_ < .001, |*d*|≥ 1.00) (Figure 5A). The distribution of consequential rates (Figure 5B) reveals that R3 has the greatest number of papers above 90% in every venue, while human reviews are broadly spread. After restricting to critical comments, the gap narrows considerably (Figure 5C–D). R3 achieves 90.0–94.8% versus 83.6–91.9% for humans, 84.3–96.4% for Gemini 3 Pro, and 81.2–90.6% for GPT-5.2.

**Figure 5:**
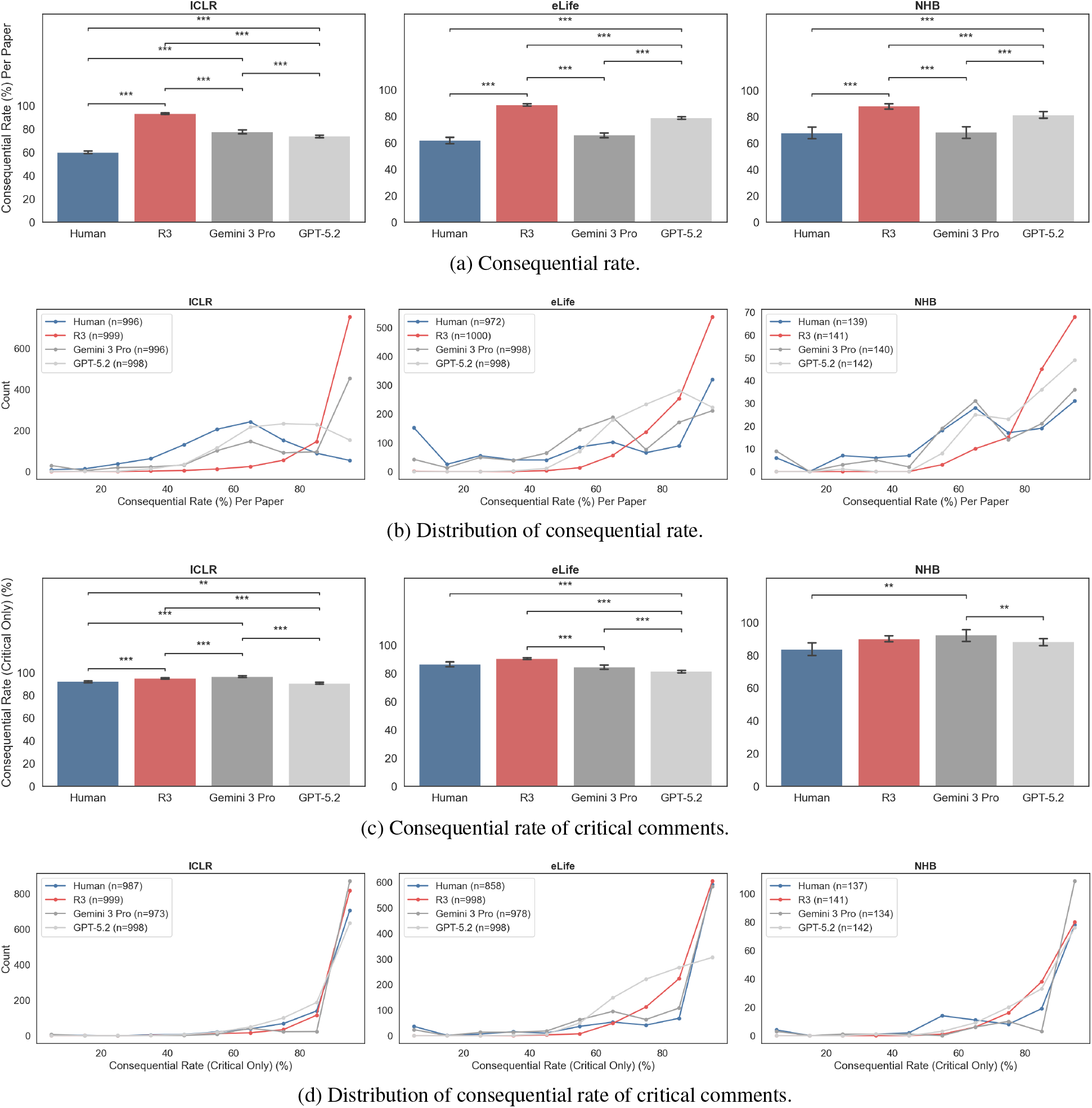
Consequential Rates. Consequential comment rate and consequential rate of critical comments by source across three venues. NHB = Nature Human Behaviour. Bars show per-paper means, error bars show 95% CIs. Brackets indicate significance (* *p* < .05, * \**p* < .01, * * \**p* < .001, Wilcoxon signed-rank, Holm–Bonferroni corrected). Cohen’s *d* reported per venue (ICLR / eLife / NHB) for pairs: Human–R3, Human–Gemini, Human–GPT, R3–Gemini, R3–GPT, Gemini–GPT (panels a, c). (a) Consequential rate: percentage of mapped comments that are consequential per paper. ICLR *n* = 992–997, eLife *n* = 970–998, NHB *n* = 137–141. *d*: −2.16, −0.78, −0.80, 0.79, 1.47, 0.18 / −1.00, −0.12, −0.62, 1.11, 0.79, −0.62 / −1.04, −0.02, −0.64, 0.93, 0.50,−0.59. (b) Distribution of consequential rate for papers with at least one mapped comment. Frequency polygon with 10% bins. (c) Consequential-critical rate: percentage of critical comments that are consequential per paper. ICLR *n* = 962–997, eLife *n* = 844–996, NHB *n* = 130–141. *d*: − 0.23, −0.34, 0.11, −0.14, 0.36, 0.45 / −0.21, 0.08, 0.28, 0.35, 0.76, 0.16 / −0.36, −0.46, −0.23, −0.17, 0.16, 0.30. (d) Distribution of consequential rate of critical comments for papers with at least one mapped critical comment. Frequency polygon with 10% bins.

### 2.6 Human Reviewers Show High Variability

We decomposed the human consequential rate distribution to individual reviewer IDs (Figure 6A). This reveals a bimodal distribution with peaks near 0% and 100% across all three venues. This pattern persists when plotting separately by acceptance decision (Figure S6). R3 ranks first most often on average consequential rate across all three venues (Figure 6B), but when restricting to critical comments (Figure 6C), human reviewers rank first on more papers than any other source in every venue despite a lower average. Together, these findings suggest that some human reviewers achieve high consequential rates that outperform AI sources, but high reviewer-to-reviewer variance may contribute to a lower human average.

**Figure 6:**
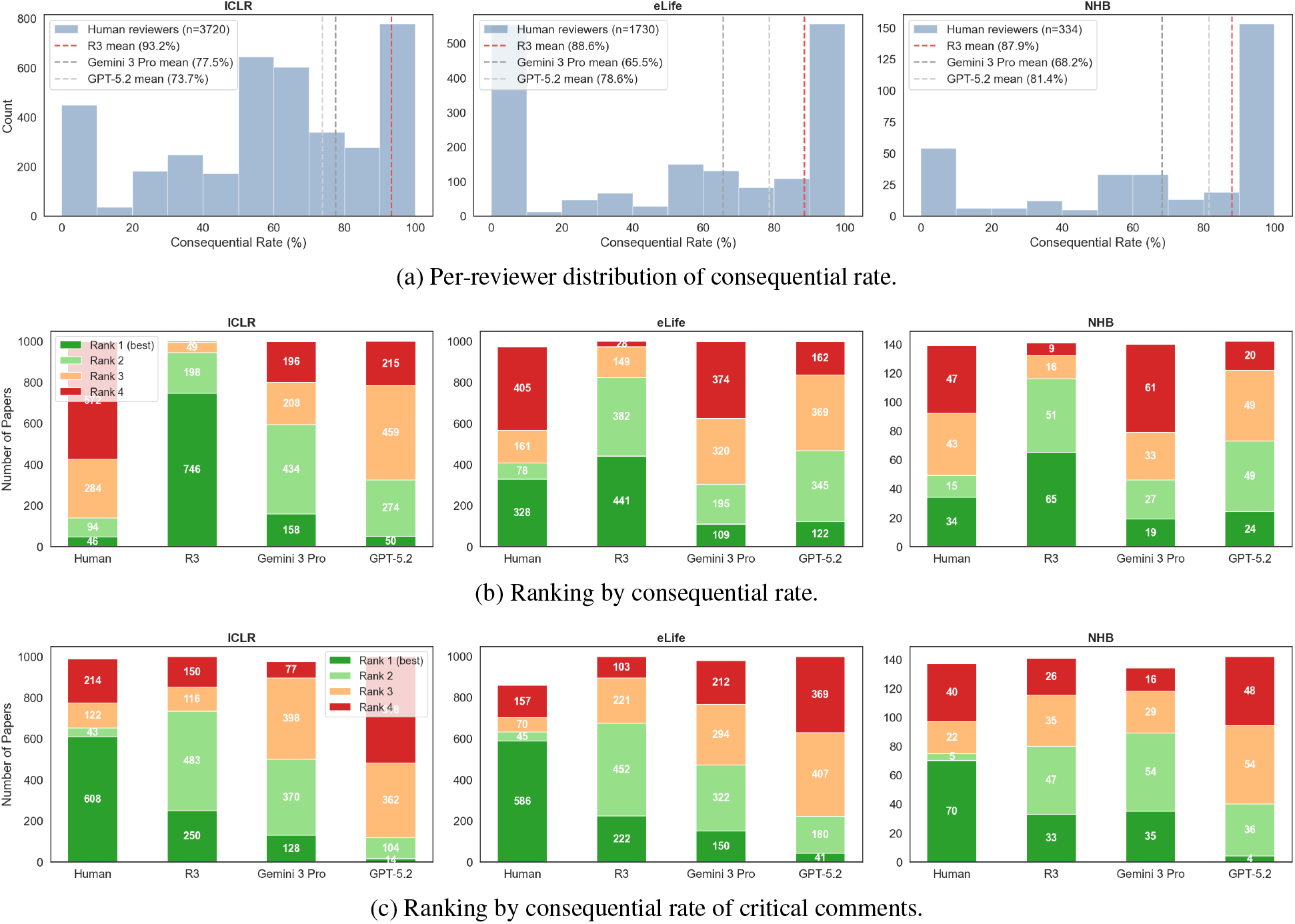
Human Reviewer Variability. Per-reviewer distribution and per-paper ranking of consequential rates across three venues. NHB = Nature Human Behaviour. (a) Per-reviewer distribution of consequential rate. Each data point is an individual reviewer’s consequential rate on a single paper. Histogram with 10% bins. (b) Per-paper ranking by consequential rate: for each paper, sources are ranked from highest to lowest consequential rate of mapped comments; stacked bars show how often each source achieves each rank. (c) Per-paper ranking by consequential rate of critical comments: for each paper, sources are ranked from highest to lowest consequential rate of critical comments; stacked bars show how often each source achieves each rank.

### 2.7 Humans Critique Contribution While AI Sources Verify

Human and AI sources provide different types of critiques (Figure 7A). Human reviewers have the highest fraction of comments related to contribution across all venues (17.5–25.2%) and the highest fraction of clarity in ICLR and Nature Human Behaviour (23.8% and 28.8%). In contrast, AI sources devote more than half of their attention to validity and sufficiency across all venues, with R3 leading in validity across all three venues (34.8–46.9%), suggesting a complementary role in technical verification.

**Figure 7:**
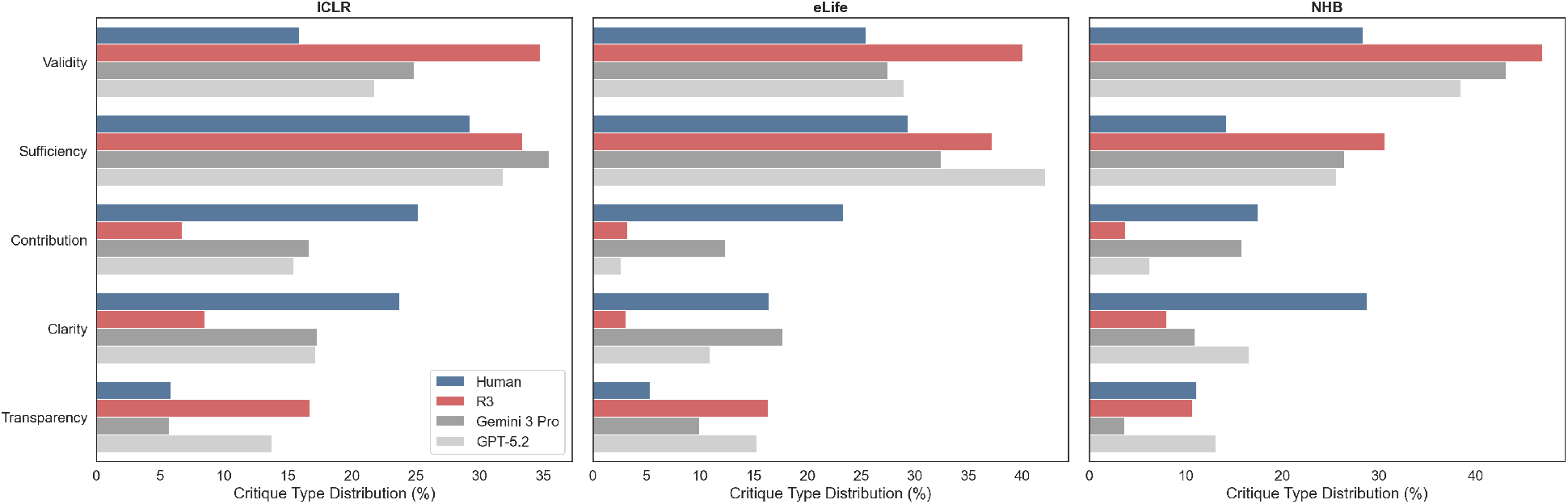
Critique Type Distribution. Critique type breakdown by source across three venues. NHB = Nature Human Behaviour. Percentage of each source’s comments assigned to each of five critique type categories. Aggregate percentages.

## 3 Discussion

As AI manuscript review tools emerge and frontier models continue to improve, benchmarks are needed to evaluate review quality in a way that is scalable and standardized. ReviewBench is an extensible and venue-agnostic framework for the direct comparison of human and AI comment structure, reviewer agreement, and critique types.

Across three disciplines, ReviewBench reveals a complementary role for AI in peer review. AI reviewers generate more structured comments that engage directly with the major claims of a paper. R3 in particular is most often consequential to those claims, suggesting it is capable of providing substantive feedback. When restricting to critical comments, per-paper rankings show human reviewers rank first on consequential rate on more individual papers than any AI source despite a lower average. The distribution of individual reviewer consequential rates similarly reveals high heterogeneity, with peaks near both 0% and 100%. This pattern persists when splitting by acceptance decision, a crude control for paper quality. Together, these data suggest that the most consequential human reviewers outperform AI sources on this metric, but a substantial fraction of reviewers near 0% brings the average down.

Combined with systematic differences in critique focus, where humans focus more on contribution and clarity, while AI sources lead on validity, sufficiency, and transparency, these results suggest that AI tools can complement human judgement. AI can reliably scale technical verification, while human judgment remains essential for evaluating contribution and shaping editorial decisions.

This analysis is limited in that it measures structural and relational properties of reviews, but not their correctness. Assessment labels are also assigned by an LLM evaluator, and although our cross-model evaluator study saw comparable source rankings across evaluators, human annotators would strengthen confidence in the results. Rule-based parsing of unstructured human reviews is also error-prone, further motivating standardized, machine-readable review formats [21]. Lastly, consequential labels apply only to claim-mapped comments, since it is by definition consequential relative to a specific claim. To mitigate penalties to sources that produce broad or editorial comments that do not map to claims, we calculate consequential rate over mapped comments only. Future iterations of ReviewBench can address these limitations as review formats become more standardized and human-annotated validation sets become available.

As manuscript submission volume continues to outpace the pool of expert reviewers, scalable benchmarks will become essential infrastructure for evaluating whether AI tools can responsibly extend reviewer capacity. ReviewBench provides a venue-agnostic pipeline for scalable assessment of human and AI reviews. Future work can extend this to support other disciplines with publicly available human reviews. As AI review tools mature, ReviewBench can serve as a living benchmark, improved by the scientific community to track progress in automated manuscript review.

## 4 Methods

ReviewBench is a publicly available [24] four-stage pipeline consisting of raw data fetchers, a layer to canonicalize raw data into papers and review comments tables, AI review generators, and a processor to extract claims from papers, assess comments, and score. The database is hosted on Cloud SQL (PostgreSQL) and is publicly accessible with read-only credentials (Database: `reviewbench`, User: `user`, Password: `public`). The benchmark is designed to be extensible to other venues or AI sources by adding a raw data fetcher and parser or generator. The input generators and processor operate on the canonical schema and require no modification. The benchmark is operable via CLI.

### 4.1 Raw Data Collection

We collected papers and human peer reviews across three venues using venue-specific fetchers available in the repository [24]. For each venue, the fetcher retrieved paper content, metadata, and peer review text and stored them as a single JSONB record per paper in the `raw_venue_data` table. No filtering was applied based on paper topic or acceptance decision.

ICLR 2025 submissions and reviews (*n* = 1,000) were retrieved via the OpenReview API v2. Nature Human Behaviour Open Access research articles with linked peer review files (*n* = 159) were scraped from the journal website. 17 articles were excluded whose peer review files did not contain structured reviewer remarks. eLife papers and peer reviews (*n* = 1,000) were loaded from a locally cloned copy of the OpenEvalProject `evals` repository [22] containing manuscript text and peer review text.

### 4.2 Canonicalization

For each venue, a venue-specific parser extracted structured paper metadata into a canonical `papers` table. Reviewer letters were split into individual comments and stored in a `comments` table. The splitting logic detects whether the text uses numbered lists, bullet lists, or paragraphs, and splits accordingly, discarding comments shorter than 20 characters. For ICLR, the Weaknesses, Questions, and Strengths fields of each structured OpenReview response were extracted and split into comments. The Summary field was excluded as it paraphrases the paper rather than evaluates it. For Nature Human Behaviour, the first round of peer review was extracted by cutting at version markers or author response boundaries, the editorial decision letter preamble was stripped, and individual reviewer letters were isolated by “Reviewer #N” headers. For eLife, peer review text was stripped of editorial boilerplate and parsed into individual reviewer letters by section dividers or “Reviewer #N” headers depending on the format. After parsing, one-time repair scripts corrected splitting artifacts (e.g. orphan section headers, editorial preambles, and multi-reviewer comments that required re-splitting). All parsing logic and repair scripts are available in the repository [24].

### 4.3 AI Reviews

R3 reviews were generated for all three venues using a multi-agent architecture with specialist reviewer agents in LLM configuration LLMcfg01.06.2026. R3 accepts either a PDF or manuscript text as input and requires no venue-specific modification.

Frontier LLM reviews were generated with Gemini 3 Pro (`gemini-3-pro-preview`) and GPT-5.2 (`gpt-5.2`) to establish a baseline for single-model LLM capability. Each paper was submitted as a PDF (ICLR, Nature Human Behaviour) or manuscript text (eLife) at temperature 0 with a minimal prompt that instructs the model to peer review the paper and return comments as a JSON array. The prompt is intentionally general and minimal so that it can be applied uniformly across venues. One eLife paper (`elife-06068`) was excluded from GPT-5.2 processing due to OpenAI content safety filtering on virology content.

### 4.4 Claim Extraction

For each paper, the manuscript PDF (ICLR, Nature Human Behaviour) or text (eLife) is supplied to `gemini-3-pro-preview` at temperature 0 with a prompt instructing it to identify 3–7 major claims. The prompt explicitly distinguishes major claims from supporting data points (e.g. specific experimental results), methodology details, novelty assertions lacking empirical support, and speculation about future work. The LLM is instructed to extract the major claims as well as the most relevant verbatim source excerpt. Claims are stored in a `claims` table keyed by paper ID.

### 4.5 Comment Assessment

For each paper, the pipeline assesses every comment from all four sources (Human, R3, Gemini 3 Pro, GPT-5.2) using three separate LLM calls to `gemini-3-pro-preview` at temperature 0, each with a dedicated prompt. The manuscript PDF or text is provided alongside comments to help resolve indirect references. To minimize redundant processing and token costs, a Gemini context cache is created containing the manuscript and reused across all three prompts.

The first prompt, comment parsing, extracts four verbatim excerpts from each review comment, or null if not explicitly stated. These are the specification (the specific issue identified), the justification (why the issue matters), the suggested remedy, and the anchor (the specific location). The second prompt, claim mapping, assigns each comment a claim ID identifying the single most relevant claim it addresses (else null). Comments mapped to claims receive a stance label (supportive, critical, or neutral), and a consequential label indicating whether the issue, if correct, could undermine the claim. The third prompt, critique typing, classifies each comment by the type of critique, drawn from a fixed taxonomy of validity, sufficiency, contribution, clarity, and transparency.

After assessment, results undergo post-processing. Claim IDs, stance labels, and critique type values are validated against the predefined sets. Invalid entries are set to `null`. Assessment results are stored in an `assessments` table keyed by comment ID and source.

### 4.6 Scoring and Visualization

The scoring step is purely computational and includes no LLM calls. All metrics are computed in Python (NumPy, pandas) from the stored assessments and claims. Figures are generated with Matplotlib, Seaborn, and statannotations. All scoring code, output CSVs, and generated figures are available in the repository [24].

Twelve metrics are calculated per paper for each source across all venues. Comments per paper is the raw count of comments. Comment length is the mean word count. Specification rate is the percentage of comments that identify a specific issue. Justification rate is the percentage that explain why the issue matters. Actionability rate is the percentage that include a suggested remedy. Anchored rate is the percentage that reference a specific location in the paper. Mapped rate is the percentage of comments that map to one of the paper’s extracted claims. Claim coverage is the percentage of a paper’s claims addressed by at least one comment. Consequential rate is the percentage of mapped comments that, if correct, could undermine the claim they address. Consequential-critical rate restricts this to comments with a critical stance. Where multiple comments map to the same claim, a dominant stance is computed as the most frequent, with ties broken by severity (critical > supportive > neutral). Claim overlap fraction is the percentage of human-addressed claims also addressed by the AI source. Stance-matched overlap fraction is the percentage of human-addressed claims also addressed by the AI source and with a matching dominant stance.

Shapiro–Wilk tests rejected normality for the majority of source–metric distributions, consistent with most metrics being bounded proportions in [0, 1]. We therefore use the Wilcoxon signed-rank test for all pairwise comparisons, with the paper as the unit of analysis, because comments within a paper are not independent. Within each venue, all four sources are assessed on the same papers. Raw *p*-values are Holm–Bonferroni corrected jointly. Because the large sample gives high power to detect trivially small differences, we report Cohen’s *d* as a measure of practical effect size. Mean differences are reported with 95% bootstrap confidence intervals.

### 4.7 Cross-Model Evaluator Study

To assess whether the evaluating LLM introduces self-preference bias, we randomly sampled 50 papers and assessed each source’s comments using GPT-5.2 with the same prompt, claims, and post-processing pipeline used by the primary Gemini evaluator. For each control source, we computed the difference between evaluator scores per paper and tested whether this difference was larger when an evaluator scored its own model’s reviews than when it scored other sources’ reviews, using a two-tailed Wilcoxon signed-rank test with Holm–Bonferroni correction across both control sources and all eight metrics. Source rank orderings were compared across all metrics.

## 5 Data and Code Availability

The ReviewBench dataset and benchmark code are publicly available at https://github.com/natalienkhalil/ ReviewBench. The database is hosted on Cloud SQL (PostgreSQL) and is publicly accessible with read-only credentials (Database: `reviewbench`, User: `user`, Password: `public`).

## 6. Acknowledgments

We ran an earlier draft of this manuscript through Reviewer3.com, which identified 1) potential evaluator self-preference bias from using Gemini as both a review source and the assessment judge, and 2) asymmetric data filtering between human and AI sources, where human reviews excluded Strengths sections for ICLR. We addressed both by adding a cross-model evaluator study, adding GPT-5.2 as a review source, and updating the human review parser to include the Strengths section.

## 7 Author Contributions

MRC conceptualized the benchmark design. NNK developed the framework and wrote the original draft. NNK and TJR conducted the investigation. All authors reviewed and edited the manuscript.

## A Supplementary Material

**Figure S1:**
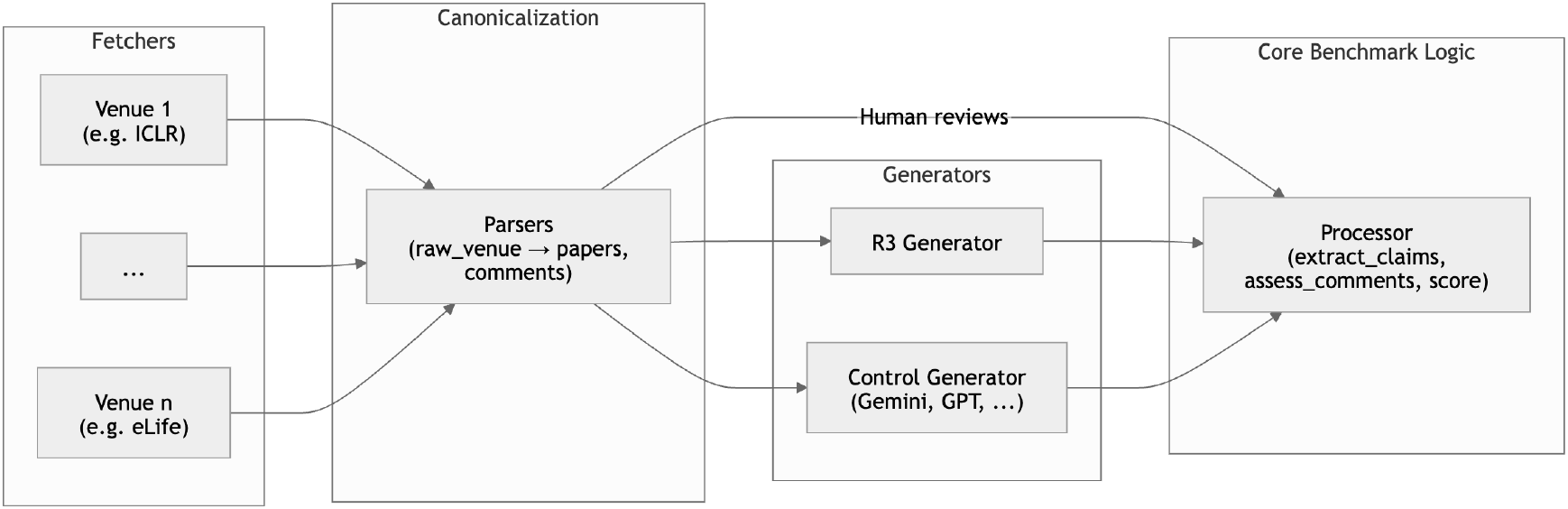
ReviewBench Design. (1) Raw data fetchers that collect papers and human reviews, (2) canonicalization into a unified schema, (3) AI review generators, and (4) processor (claim extraction, comment assessment, and scoring).

**Figure S2:**
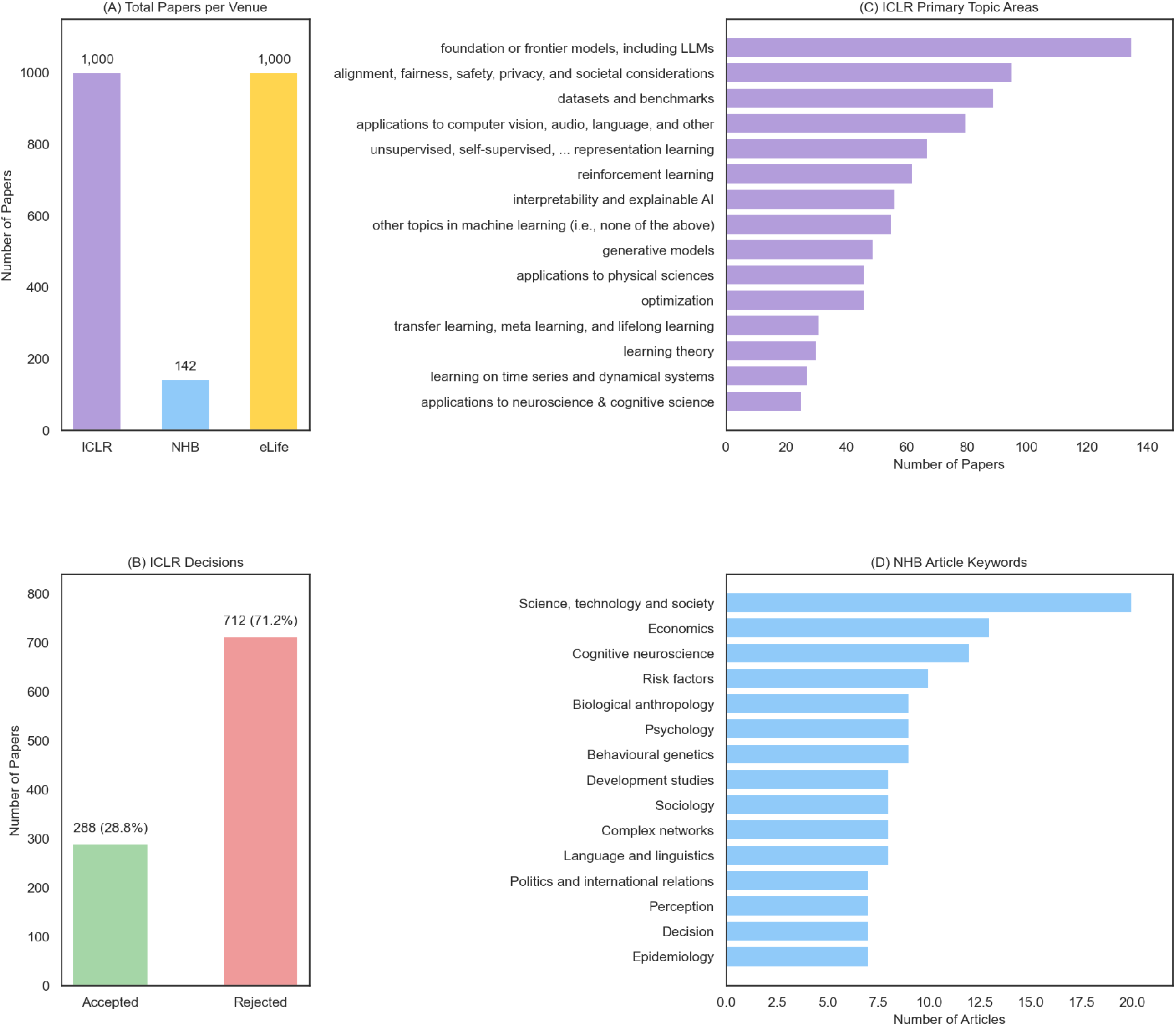
Dataset Overview Across Venues. (A) Total papers per venue. ICLR (*n* = 1,000), Nature Human Behaviour (*n* = 142), and eLife (*n* = 1,000). (B) ICLR acceptance decisions. 288 accepted (28.8%) and 712 rejected (71.2%). (C) Distribution of ICLR papers by primary topic area (top 15). (D) Distribution of Nature Human Behaviour articles by keyword (top 15).

**Figure S3:**
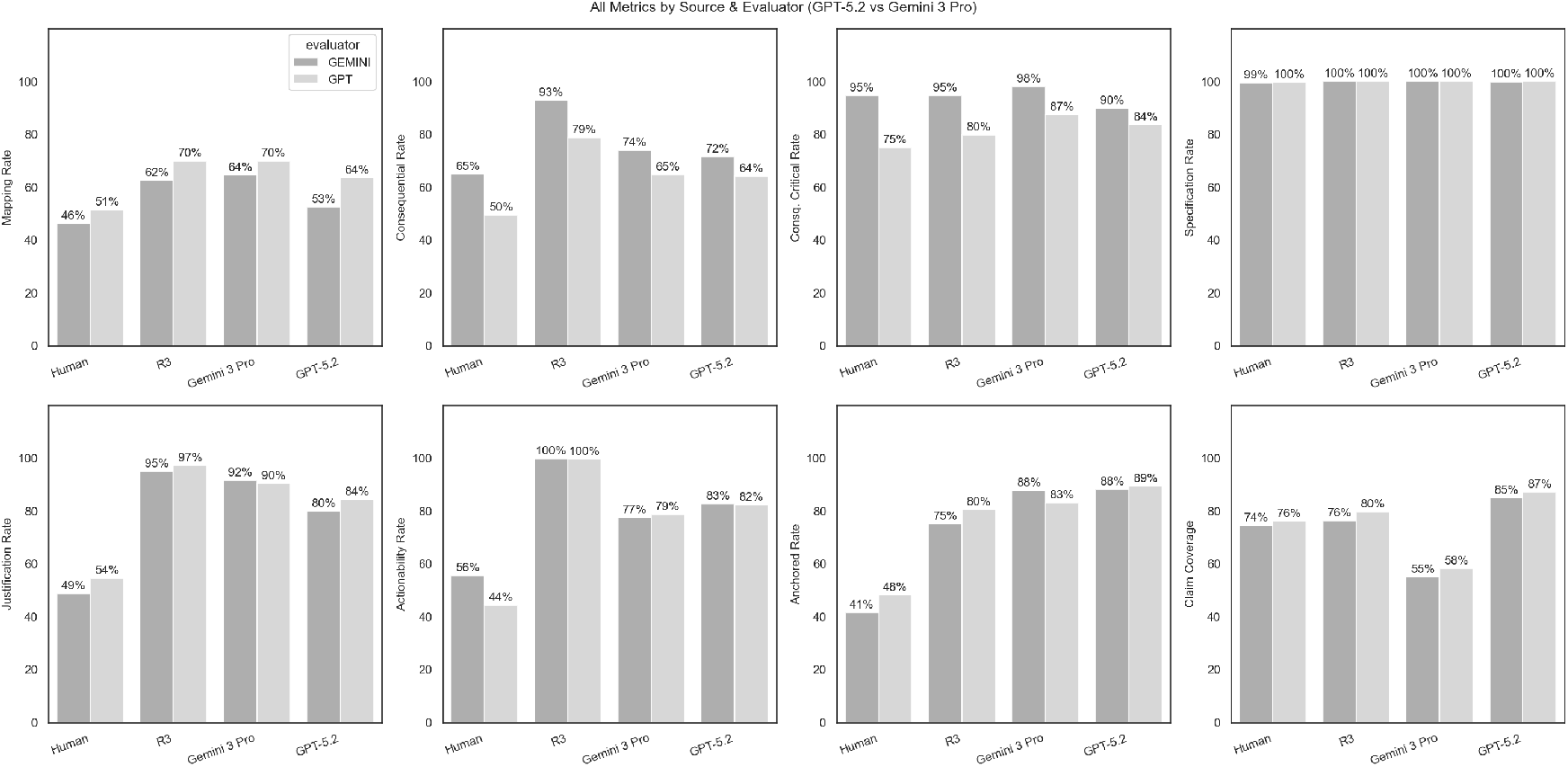
Cross-Model Evaluator Study: Metrics by Evaluator. Per-paper metric values when assessed by Gemini 3 Pro vs. GPT-5.2 evaluators across all eight metrics computed in the study. Each panel shows paired evaluator scores for all four sources. *n* = 50 randomly sampled papers.

**Figure S4:**
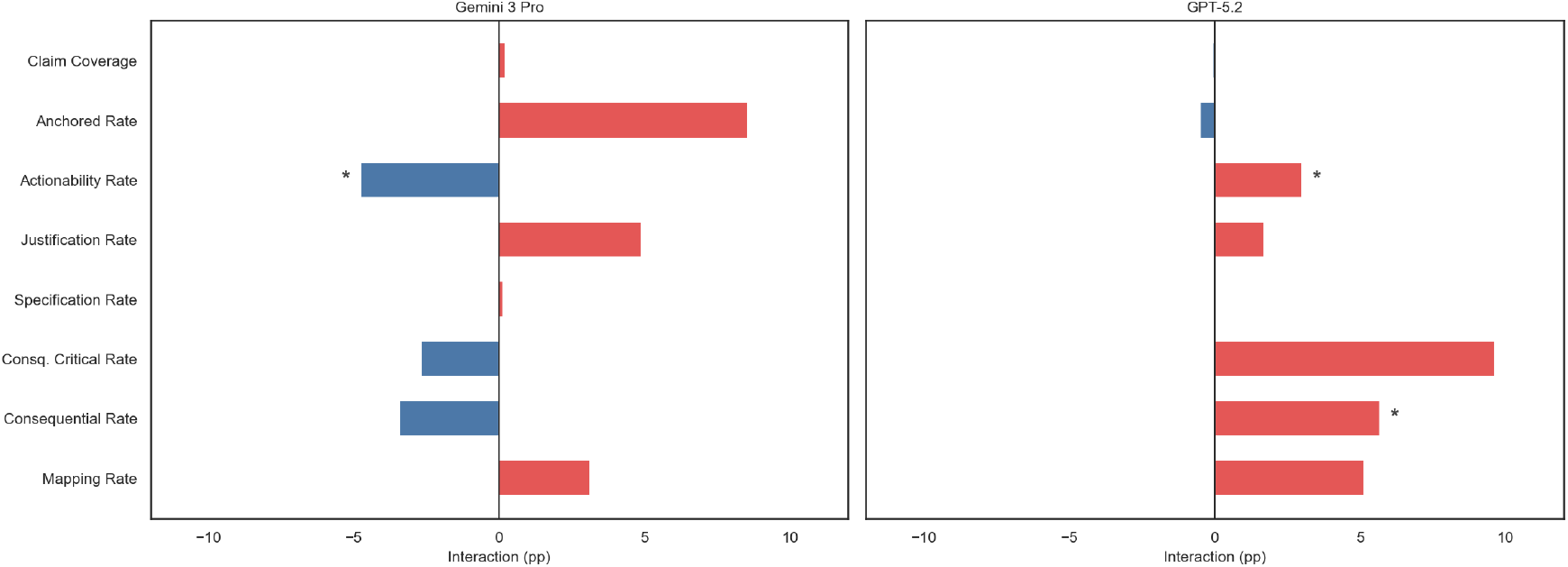
Cross-Model Evaluator Study: Self-Favoritism Analysis. Mean interaction effect per metric from a mixed-effects model with Wilcoxon signed-rank test. Positive values indicate self-preference, where evaluator scores its own model higher. * indicates *p* < .05 (Holm–Bonferroni corrected). *n* = 50 papers.

**Figure S5:**
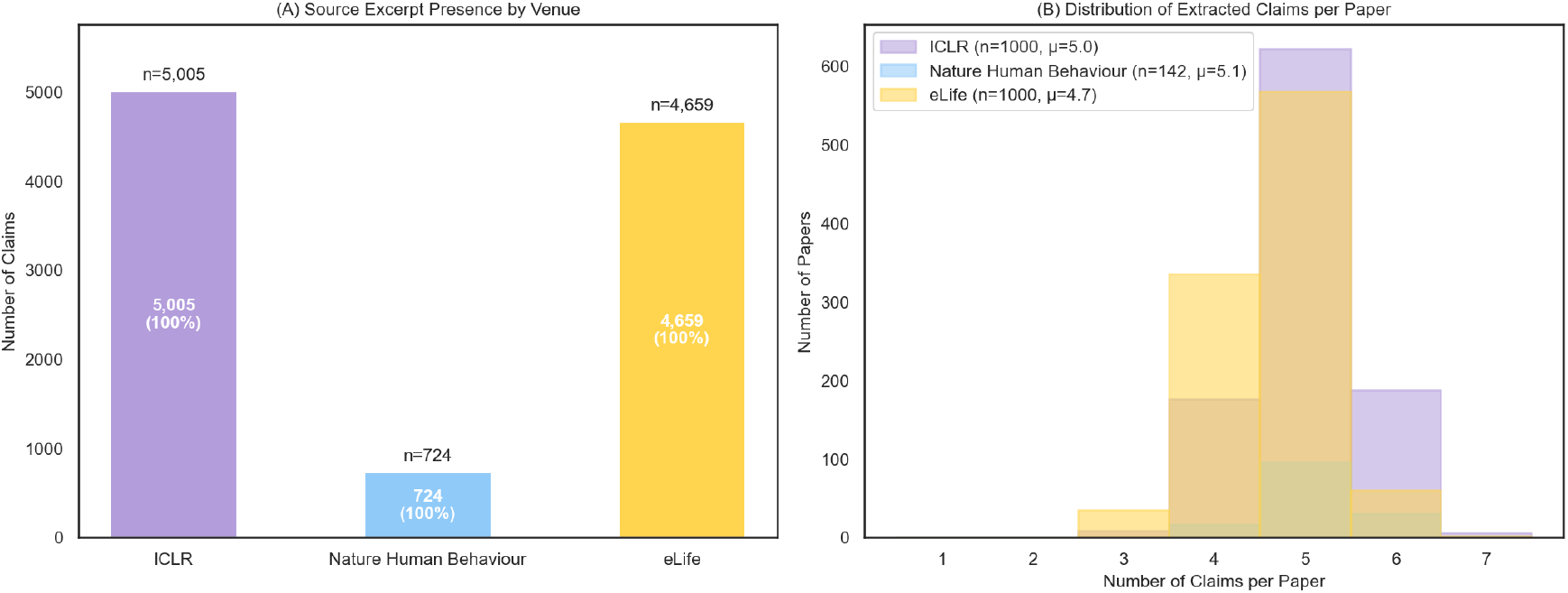
Claim Extraction Overview Across Venues. (A) Source excerpt presence by venue. Stacked bars show the number of extracted claims with and without a verbatim source excerpt from the paper. ICLR (*n* = 5,005 claims), Nature Human Behaviour (*n* = 724), and eLife (*n* = 4,659). (B) Distribution of major claims extracted per paper across all three venues. Mean claims per paper: ICLR = 5.0, Nature Human Behaviour = 5.1, eLife = 4.7.

**Figure S6:**
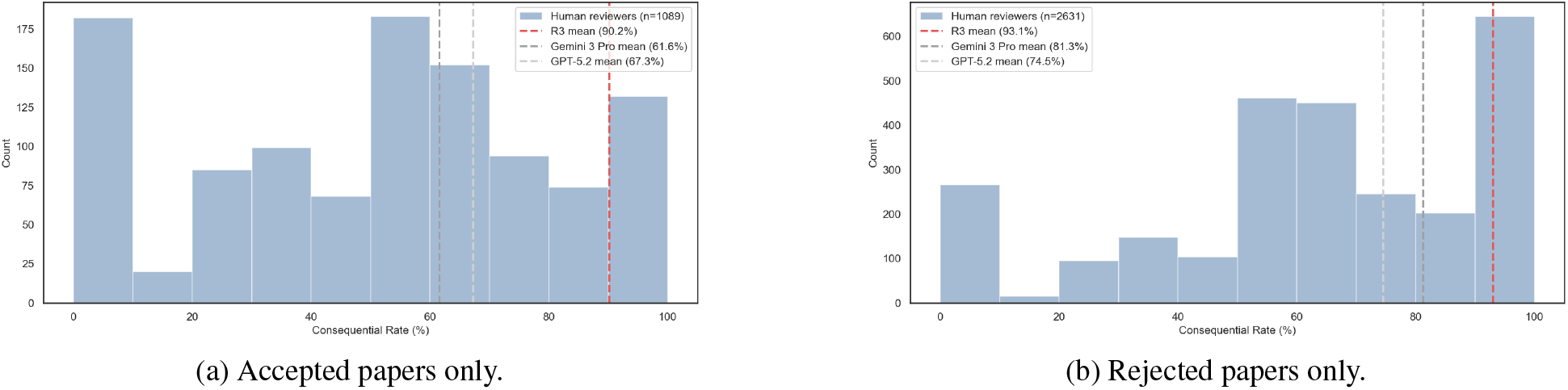
Reviewer Consequential Rate Distribution by Submission Decision. Distribution of consequential rate split by submission decision for ICLR 2025 submissions. (a) Accepted papers only (*n* = 1,089 reviewer–paper pairs, mean = 49.5%). (b) Rejected papers only (*n* = 2,631 reviewer–paper pairs, mean = 62.2%). The bimodal pattern with peaks near 0% and 100% persists in both subsets, indicating that reviewer-to-reviewer variance is not solely driven by paper quality.

**Table S1:** Source Rankings by Metric Across Evaluators. For each of eight scoring metrics, sources are ranked by mean value under the Gemini 3 Pro and GPT-5.2 evaluators independently. “Match” indicates whether both evaluators produce the same source ranking. *n* = 50 randomly sampled papers.

| **Metric** | **Gemini Ranking** | **GPT Ranking** | **Match** |
| --- | --- | --- | --- |
| Mapped Rate | Gemini 3 Pro > R3 > GPT-5.2 > Human | Gemini 3 Pro > R3 > GPT-5.2 > Human | Yes |
| Consequential Rate | R3 > Gemini 3 Pro > GPT-5.2 > Human | R3 > Gemini 3 Pro > GPT-5.2 > Human | Yes |
| Consequential Rate (Critical Only) | Gemini 3 Pro > Human > R3 > GPT-5.2 | Gemini 3 Pro > GPT-5.2 > R3 > Human | No |
| Specification Rate | Gemini 3 Pro = R3 > GPT-5.2 > Human | Gemini 3 Pro = R3 = GPT-5.2 > Human | Yes |
| Justification Rate | R3 > Gemini 3 Pro > GPT-5.2 > Human | R3 > Gemini 3 Pro > GPT-5.2 > Human | Yes |
| Actionability Rate | R3 > GPT-5.2 > Gemini 3 Pro > Human | R3 > GPT-5.2 > Gemini 3 Pro > Human | Yes |
| Anchored Rate | GPT-5.2 > Gemini 3 Pro > R3 > Human | GPT-5.2 > Gemini 3 Pro > R3 > Human | Yes |
| Claim Coverage | GPT-5.2 > R3 > Human > Gemini 3 Pro | GPT-5.2 > R3 > Human > Gemini 3 Pro | Yes |

**Table S2:** Example Comments by Stance Category. Representative comments classified as supportive, critical, or neutral for human and AI sources. One example per category per source is shown.

| Stance | Source | Venue | Paper | Comment |
| --- | --- | --- | --- | --- |
| Supportive | Human | ICLR | *FCoReBench: Can Large Language Models Solve Challenging First-Order Combinatorial Reasoning Problems?* | “The proposed SymPro-LLM can work with different instances from the same first order combinatorial optimization problem without the need to re-evaluate using LLMs.” |
| Supportive | R3 | eLife | *The human ARF tumor suppressor senses blastema activity and suppresses epimorphic tissue regeneration* | “While the authors correctly note the absence of a direct ARF ortholog in highly regenerative vertebrates, the discussion should address whether these species utilize alternative, non-homologous mechanisms to sense Rb-pathway hyperphosphorylation. Discussing how organisms like zebrafish or urodeles manage oncogenic E2F signals in the absence of ARF would provide a more complete evolutionary context for why the blastema is permitted to proliferate without triggering alternative tumor suppressor responses.” |
| Critical | Human | NHB | *Consonant lengthening marks the beginning of words across a diverse sample of languages* | “The discussion of potential phonotactic cues to word segmentation ([language-]specific complex clusters’) is rather convoluted and could be shortened: some languages have effective consonant clustering cues and some do not.” |
| Critical | R3 | eLife | *The AFF4 scaffold binds human P-TEFb adjacent to HIV Tat* | “The use of a thermodynamic cycle to infer Tat’s affinity for the AFF4-P-TEFb complex is an elegant workaround for Tat’s in vitro instability. However, this calculation assumes that the structural integrity and binding interfaces of P-TEFb remain identical whether Tat or AFF4 binds first. The authors should briefly discuss the limits of this assumption, particularly whether allosteric changes induced by one binding partner could cause the actual cooperative binding affinity in vivo to deviate from the calculated in vitro values.” |
| Neutral | Human | ICLR | *Double Check My Desired Return: Transformer with Value Validation for Offline RL* | “It will be exciting if Doctor is able to show clear improvements in online fine-tuning experiments on AntMaze or Kitchen or Adroit tasks, where offline-to-online methods (e.g. Cal-QL) have showed significant improvements. Q3. Other than Q1 (final performance) and Q2 (online fine-tuning), can you share your ideas on why having better return alignment is beneficial, if there is any?” |
| Neutral | R3 | NHB | *Human ventromedial prefrontal cortex is necessary for prosocial motivation* | “The reported dissociation between medial (area 14) and lateral (area 13) vmPFC damage on prosociality is a major finding, yet it is primarily contextualized using macaque electrophysiology. The authors should expand the discussion to integrate existing human functional neuroimaging literature on self versus other valuation, specifically addressing whether fMRI studies of prosocial giving or vicarious reward in healthy humans support a similar medial-lateral gradient.” |

**Table S3:** Example Consequential Comments. A consequential comment is defined as one that could undermine the mapped claim. One example per source is shown.

| Source | Venue | Paper | Comment |
| --- | --- | --- | --- |
| Human | ICLR | *Overcoming Slow Decision Frequencies in Continuous Control: Model-Based Sequence Reinforcement Learning for Model-Free Control* | “In line 150 the claim that online planning is not feasible in robotics or the brain is made. In my experience this is not true. How did the authors come to this conclusion?” |
| R3 | eLife | *fMRI reveals neural activity overlap between adult and infant pain* | “Stimulus comparability and interpretation: Adults received up to 512 mN on dorsum of foot, infants up to 128 mN on heel. Differences in site (heel vs dorsum), tissue properties, receptor density, and force range complicate inference that infants are ‘more sensitive’ based on % active voxels. Consider reframing as ‘different stimulus–response functions’ and add justification or calibration (e.g., mechanical indentation depth, contact area, or perceptual equivalence in adults at heel).” |
| Gemini 3 Pro | eLife | *Registered report: Biomechanical remodeling of the microenvironment by stromal caveolin-1 favors tumor invasion and metastasis* | “In Protocol 3, the experimental timeline is set to 70 days, but standard euthanasia criteria (e.g., tumor exceeding 20 mm) are also listed. If the primary tumors grow faster than anticipated and mice must be euthanized before day 70, how will this affect the ex vivo metastasis quantification? The authors should clarify if metastasis will be assessed at the time of early euthanasia and how varying experimental durations will be handled in the statistical analysis.” |
| GPT-5.2 | ICLR | *A General Feature Attribution Framework under a Black-box Setting* | “Minor issue (text experiment deletion types): The paper uses two deletion operations (embedding reset vs token removal, Sec. 5.2 p.8) and notes IG uses zero-embedding baseline. For fairness, did GEFA use the same baseline as the deletion operation being evaluated (empty token vs zero embedding)? Please clarify the baseline used during explanation for each deletion evaluation; otherwise the comparison may conflate explanation method with absence definition.” |

